# Environmental filtering across seasons and host-associated selection shape the gut microbiome of sympatric European *Sepsis* dung flies

**DOI:** 10.64898/2026.06.24.734284

**Authors:** Martin Kapun, Jeannine Roy, Wolf U. Blanckenhorn

## Abstract

Animal microbiomes are shaped by both environmental exposure and host-associated filtering, but the relative importance of these processes remains poorly understood. Dung-associated insects provide an ideal model because they develop and feed in highly dynamic microbial environments. We investigated the gut microbiomes of six sympatric dung fly species of the genus *Sepsis* (Diptera: Sepsidae) and compared them with microbial communities in cow dung throughout a growing season in Switzerland. Using full-length 16S rRNA gene sequencing (PacBio), we characterized bacterial communities from 74 fly and 15 dung samples.

Seasonal variation was the strongest predictor of microbiome composition, whereas host species exerted weaker effects that persisted after removing dung-associated taxa, indicating that gut communities are not merely passive reflections of environmental exposure. Only few gut microbiome reads were attributable to dung-associated taxa, and environmental overlap differed among fly species rather than season. A highly non-random core microbiome persisted across all six species: 36 bacterial genera (of 469) were shared by all hosts at ∼119-fold enrichment above random expectation and remained after removing dung-associated taxa. These findings support a two-layer model of microbiome assembly, in which seasonal environmental variation determines microbial availability while host-specific processes selectively retain a subset of taxa.

## Introduction

Animal-associated microbial communities play a fundamental role in physiological, ecological and evolutionary aspects of their host organisms. Such factors include nutrition, development, immunity, behavior, and ecological adaptation (Shapira, 2016; Macke *et al*., 2017; Douglas, 2018; Koskella and Bergelson, 2020). Over the past decades, advances in high-throughput sequencing have allowed characterization of microbial community composition and dynamics in unprecedented detail, revealing that microbiomes are integral components of animal biology rather than passive collections of commensal organisms (Engel and Moran, 2013; Douglas, 2015). At the same time, it has become increasingly clear that the ecological and evolutionary processes governing microbiome assembly differ markedly among taxa. Whereas some animals actively maintain and influence highly specialized and evolutionarily stable microbial symbioses (see, for example, O’Brien *et al*., 2021), others acquire much of their microbiota from the surrounding environment and exhibit substantial temporal variation in community composition, as found, for example, in mosquitos (Ravenscraft and Coon, 2026) or rodents (Fenn *et al*., 2023). Understanding the balance between environmental acquisition and host-associated filtering therefore remains a central challenge in microbiome ecology (Bordenstein and Theis, 2015; Douglas and Werren, 2016; Theis *et al*., 2016; Foster *et al*., 2017; Simon *et al*., 2019; Vliet and Doebeli, 2019).

Insects provide particularly valuable systems for investigating microbiome assembly because they occupy an extraordinary diversity of ecological niches and exhibit remarkable variation in the degree of microbial specialization (Engel and Moran, 2013; Yun *et al*., 2014; Douglas, 2015). In some groups, such as aphids or tsetse flies, obligate bacterial symbionts have co-evolved intimate associations with their hosts to provide essential functions (Provorov and Vorob’ev, 2012; Kolasa *et al*., 2019; Attardo *et al*., 2020). In contrast, many holometabolous insects harbor rather flexible microbial communities that are acquired repeatedly from the environment and may change substantially throughout development or across ecological contexts (Hammer *et al*., 2017, 2019). A key challenge in this context is distinguishing environmentally derived microbes - those passively ingested from the surroundings - from stable host-associated communities that persist across environments and individuals (see, for example, Ludington, 2024). This distinction is critical for understanding whether and which gut microbiomes and specific microbes have ecological or evolutionary relevance for their hosts.

Dung-associated insects are especially informative for studying these questions because they develop and live within one of the most microbially rich and dynamic habitats in terrestrial ecosystems (Shukla *et al*., 2016; Suárez-Moo *et al*., 2020; Ebert *et al*., 2021). Vertebrate dung supports dense and rapidly changing microbial communities that drive decomposition, nutrient cycling, and energy transfer through food webs (Girija *et al*., 2013; Holter, 2016). Fresh dung is rapidly colonized by bacteria and fungi that transform complex organic substrates into forms accessible to a diverse assemblage of invertebrates (Yamada *et al*., 2007; Geiger, 2010; Pecenka and Lundgren, 2018; Zhang *et al*., 2025). These microbial communities change dramatically over time as dung ages, moisture content decreases, and competitive interactions alter community structure (Li *et al*., 2020; Palumbo *et al*., 2021). Insects exploiting dung are therefore exposed to continuously shifting and diverse environmental microbial pools across a seasonal gradient, which allows to disentangle host-associated from environmentally acquired gut microbiota provided that environmental communities are sampled in parallel.

Coprophagous insects play key ecological roles within these systems. Through feeding, burrowing, and dispersal activities they influence decomposition rates, nutrient recycling, parasite suppression, and the redistribution of microorganisms among dung pats (Yamada *et al*., 2007; Pecenka and Lundgren, 2018). While dung beetles have received considerable attention because of their well-known ecosystem services (Nichols *et al*., 2008; Hanski and Cambefort, 2014; Holter, 2016), dung flies often numerically dominate dung-associated insect communities in more temperate climates and thus contribute substantially to decomposition processes (Floate, 2011, 2023; Rohner *et al*., 2015; Sigsgaard *et al*., 2021). Despite their ecological importance and intimate association with a microbially dynamic substrate, relatively little is known about the microbiomes of dung-associated flies or the processes shaping microbial community assembly within their digestive tracts.

Species of the genus *Sepsis* (Diptera: Sepsidae) are among the most common and widespread dung-associated flies throughout Europe (Pont, 1986; Pont and Meier, 2002; Rohner *et al*., 2015; Rohner, Haenni, *et al*., 2019). Multiple species frequently co-occur within the same dung pats and exploit similar substrates throughout their larval development. Nevertheless, species exhibit differences in phenology, body size, life-history traits, behavioural ecology and dung preference (Blanckenhorn *et al*., 1999, 2020, 2021; Rohner *et al*., 2015; Rohner, Haenni, *et al*., 2019; Rohner, Roy, *et al*., 2019). These differences raise the possibility that sympatric species partition resources in subtle ways that facilitate coexistence despite strong apparent overlap in habitat use (Laux *et al*., 2019). Because larvae feed directly on dung-associated microorganisms and decomposing organic matter, microbial communities may represent an underappreciated axis of ecological differentiation, particularly if different host species selectively retain distinct subsets of environmental microorganisms.

Microbiome variation among sympatric species could arise through several non-exclusive mechanisms. First, host species may differ in gut physiology, immune function, or digestive environment, leading to selective retention of different subsets of the available environmental microbiota (e.g., Phalnikar *et al*., 2018; Li *et al*., 2026). Second, species may exploit distinct microhabitats or feeding substrates within dung pats, generating differential exposure to microbial communities even within the same dung pat (Su *et al*., 2024). Third, seasonal variation in environmental microbial communities may impose a shared temporal signal on all co-occurring species simultaneously, generating turnover that masks or interacts with host-specific effects (Akorli *et al*., 2016; Naveed *et al*., 2024). Fourth, phylogenetic relatedness among host species may influence microbiome composition through long-term host–microbe associations. Closely related hosts may retain similar microbial lineages as a consequence of shared ancestry, co-evolutionary processes, and conserved host traits, resulting in greater microbiome similarity among related species than among more distantly related taxa (Martinson *et al*., 2017; Mallott and Amato, 2021). Disentangling these processes requires simultaneous characterization of both host-associated and environmental microbial communities across multiple host species and time points.

Although microbiome studies have become increasingly common across insects, virtually nothing is known about microbiome assembly in sepsid flies. Consequently, it remains unclear whether co-occurring *Sepsis* species harbor distinct gut microbiomes, whether seasonal environmental variation drives community composition, to what extent fly gut communities simply reflect the microbiota of their dung substrate, and whether any microbiome taxa are genuinely host-associated rather than transiently acquired. Addressing these questions requires integrating environmental and host-associated sampling within a rigorous ecological framework.

In the present study, we conducted the first comprehensive assessment of gut microbiome variation among sympatric *Sepsis* species. Six dung-fly species and their associated cow-dung communities were sampled across an entire growing season, and bacterial community composition was characterized using full-length 16S rRNA amplicon sequencing. Unique amplicon sequence variants (ASVs) were identified and taxonomically classified using the SILVA 138.2 reference database. To explicitly distinguish environmentally acquired from host-associated microbiome components, we identified gut ASVs shared with dung communities and integrated this information with redundancy analyses, indicator-species analyses, and multi-species taxon-sharing tests.

We tested four primary hypotheses. First, we predicted that host species would differ in gut microbiome composition reflecting species-specific filtering of environmental microorganisms. Second, we predicted that seasonal variation would exert a strong and consistent influence on microbiome composition, because dung communities undergo rapid temporal turnover throughout the growing season. Third, we predicted that the degree of environmental (dung-derived) contamination in gut communities would differ among host species reflecting differences in habitat use or filtering capacity. Fourth, we predicted that a set of bacterial genera would be shared among host species at frequencies exceeding random expectation, constituting a non-random core microbiome that persists despite environmental turnover. Together, these analyses provide new insights into microbiome assembly in natural insect populations and contribute to a broader understanding of how host identity and environmental variation interact to structure microbial communities in decomposer ecosystems.

## Material and Methods

### Study System and Sampling Design

Adult dung flies of the genus *Sepsis* (Diptera: Sepsidae) and cow-dung samples were collected from a dairy pasture at Ziegelhütte/Schwamendingen near Zürich, Switzerland throughout the 2020 growing season. Sampling was conducted monthly between April and September, covering the period of peak sepsid activity and seasonal turnover in dung-associated insect communities.

During each sampling event, we collected adult flies using standardized sweep-net sampling. A total of 600 sweeps were performed at randomly selected positions throughout the pasture during each visit. In parallel, we sampled three fresh cow-dung pats per sampling event to characterize the environmental microbial community (see Table S1).

Approximately 25 ml of dung was collected from each pat and transferred into sterile Falcon tubes. All samples were transported to the laboratory, transferred into 96% ethanol to wash away microbes on the surface of the flies and subsequently frozen at −20°C within two hours of collection.

We subsequently identified male sepsid flies to species level using diagnostic morphological characters under a stereomicroscope (Pont and Meier, 2002). To maximize taxonomic representation and temporal replication, six focal species were further selected for microbiome analyses: *Sepsis cynipsea*, *S. duplicata*, *S. flavimana*, *S. neocynipsea*, *S. punctum,* and *S. thoracica*. Samples were pooled by species and collection date and preserved in 96% ethanol until DNA extraction. In total, 91 fly and dung samples were included in our subsequent analyses (see Table S2).

Because male *S. thoracica* exhibit pronounced body-size and colour polymorphism (Busso *et al*., 2017; Busso and Blanckenhorn, 2018b, 2018a; Blanckenhorn *et al*., 2020; Gourgoulianni *et al*., 2023), additional within-species sampling was performed. We selected the two largest and two smallest males available from each collection period for downstream analyses whenever possible. To this end, we mounted the femora on microscope slides using Euparal mounting medium, photographed at 40-fold magnification, and estimated body size based on the digitally measured femur lengths of the right foreleg.

### DNA Extraction

Microbial DNA was extracted using the ZymoBIOMICS DNA Microprep Kit (Zymo Research), which is optimized for efficient recovery of microbial DNA from both Gram-positive and Gram-negative bacteria (Burjanivova *et al*., 2026). Prior to extraction, we carefully air-dried ethanol-preserved flies for approximately five minutes to remove residual ethanol. We then homogenized individual flies in 200 µl lysis buffer using sterile plastic pestles and transferred the homogenates into bead-beating tubes supplied with the extraction kit, supplemented with an additional 550 µl lysis buffer, and disrupted mechanically for five minutes at maximum speed. DNA extraction was subsequently completed according to the manufacturer’s protocol.

Dung samples were processed using the same extraction kit. A small amount of frozen dung was transferred directly into bead-beating tubes and extracted without protocol modifications. Subsequently, DNA concentrations were quantified spectrophotometrically and stored at −20°C until amplification.

### Amplification of Full-Length 16S rRNA Genes

Bacterial communities were characterized using full-length amplification of the 16S rRNA gene. PCR amplification followed the PacBio protocol for full-length bacterial 16S sequencing (PacBio, 2022). Amplification employed combinations of barcoded 27F (5′-AGRGTTYGATYMTGGCTCAG-3′) and 1492R (5′-RGYTACCTTGTTACGACTT-3′) primers.

Forward and reverse primers contained unique barcode combinations, allowing multiplexing of all samples within a single sequencing run. Primers additionally included 5′ phosphate modifications and buffer sequences required for downstream library preparation.

PCR reactions were performed using KAPA HiFi HotStart ReadyMix (KAPA Biosystems), a proofreading polymerase optimized for long-range amplification. The target amplicon length was approximately 1,600 bp. Amplification success was assessed by electrophoresis on 1.5% agarose gels. PCR products were subsequently grouped into concentration classes based on band intensity and DNA concentration estimates. To compensate for variation in amplification yield, samples were pooled in unequal volumes according to concentration class to generate an approximately equimolar amplicon pool. Final DNA concentration was quantified using a Qubit High Sensitivity assay (Thermo Fisher Scientific), and fragment size distributions were verified using an Agilent Bioanalyzer.

### Library Preparation and PacBio Sequencing

Library preparation and sequencing were conducted at the Functional Genomics Center Zürich (FGCZ). Amplicon pools were first subjected to DNA damage repair and polishing before ligation of SMRTbell adapters using the SMRTbell Express Template Preparation Kit 2.0 (Pacific Biosciences). Sequencing complexes were generated using the Sequel II Binding Kit 2.1 and sequenced on a PacBio Sequel II instrument using a single SMRT Cell 8M and a 15-hour movie collection time.

Raw sequence data were processed using PacBio SMRT Link software to generate circular consensus sequences (CCS), demultiplex reads based on barcode combinations, and perform initial quality filtering.

### Bioinformatic Processing

Raw CCS reads were converted to FASTQ format using samtools (v.1.18, Li *et al*., 2009) and quality assessed using LongQC (v.1.20, Fukasawa *et al*., 2020). Read-length distributions, quality scores, and coverage profiles were inspected to identify potential sequencing artefacts and non-target amplicons. We then processed sequences using DADA2 as implemented in QIIME2 (v.2026.4, Estaki *et al*., 2020). Primer sequences were removed and reads lacking complete primer sequences were discarded. Moreover, we excluded reads outside the expected full-length 16S size range. DADA2 was then used to perform dereplication, error correction, amplicon sequence variant (ASV) inference, and chimera removal.

To identify non-bacterial contaminants, all ASVs were exported and searched against a local copy of the NCBI nucleotide database using BLASTN (v.2.12, Altschul *et al*., 1990; Camacho *et al*., 2009). This screening revealed the presence of plant plastid sequences and eukaryotic ribosomal DNA, including host-derived 18S sequences specific to sepsid flies. We removed these ASVs from subsequent analyzes and further excluded samples containing fewer than 1,000 reads after filtering. Rarefaction analyses indicated that diversity estimates reached asymptotes at approximately 1,000-5,000 reads for most samples. To standardize sampling effort across libraries, the dataset was rarefied to 1,392 reads per sample, corresponding to the minimum sequencing depth among retained samples.

### Taxonomic Classification

We performed taxonomic assignments using the SILVA v138.2 database (Quast *et al*., 2013) and classified consensus sequences with the VSEARCH consensus-classifier algorithm implemented in QIIME2. Taxa were assigned to the lowest possible taxonomic level and summarized at multiple hierarchical levels including phylum, class, order, family, genus, and ASV. To display the microbial diversity among samples, we generated relative abundance plots in R (R Core Team, 2019) using ggplot2 (Wickham *et al*., 2019). Only taxa exceeding 5% relative abundance in at least one sample were displayed individually, while remaining taxa were grouped as low-abundance categories.

### Alpha Diversity Analyses

Within-sample diversity was quantified using multiple indices calculated from rarefied ASV tables using the *core-metrics-phylogenetic* and *diversity alpha* functions of QIIME2. The Shannon index characterized overall diversity accounting for richness and rare taxa. The Simpson index and Pielou’s evenness (J’) index were used to assess community evenness and dominance structure. Chao1 and observed-feature richness indices were included to capture taxon richness. Effects of sampling month (“Season”), host species (“Species”), and their interaction were evaluated using two-way analysis of variance (ANOVA) in R.

### Beta Diversity Analyses

Differences in microbial community composition among samples were quantified using Bray-Curtis dissimilarity and weighted UniFrac distances using the *beta-group-significance* function of QIIME2. Bray-Curtis dissimilarities capture differences in taxon abundance and presence-absence among communities, respectively, whereas UniFrac distances additionally incorporate phylogenetic relationships among taxa. For UniFrac analyses, ASVs were aligned using MAFFT and a phylogenetic tree was reconstructed with FastTree using the implementations of both programs in QIIME2. Subsequently, we employed principal coordinates analyses (PCoA) using the *pcoa* function in QIIME2 to visualize community variation and assessed statistical significance of differences among seasons and species using permutational multivariate analysis of variance (PERMANOVA) with 999 permutations. Pairwise *P*-values were adjusted for multiple comparisons using the Benjamini-Hochberg false-discovery rate procedure.

### Redundancy Analysis and Indicator Taxa

To identify ecological drivers of microbiome composition, redundancy analyses (RDA) were performed using the vegan package in R (Dixon, 2003). Rare taxa were removed by retaining only ASVs or aggregated taxa with at least 100 reads across the dataset. Count data were Hellinger transformed prior to analysis to reduce the influence of differences in total abundance among samples and to make the data suitable for Euclidean-based multivariate analyses. We included species identity and sampling month as explanatory variables and assessed statistical significance using permutation tests with 999 permutations, testing marginal effects of each predictor using the Type III (marginal) approach. Forward and backward model selection was performed using the *ordiR2step* and *ordistep* functions to identify the minimum adequate model. RDA was performed at all taxonomic levels (phylum to species and ASV) and repeated for the full dataset, the dung-associated taxa subset, and the dung-filtered dataset.

To identify taxa associated with particular host species or sampling periods, indicator-species analyses were performed using the multipatt function implemented in the indicspecies package in R (Cáceres and Legendre, 2009); we assessed statistical significance using 1000 permutations.

### Environmental Dung Microbiome and Core Microbiome Analyses

Dung samples were processed using the same bioinformatic pipeline as fly samples. Because sequencing depth was substantially lower in dung libraries following quality filtering, we pooled all dung samples to generate a comprehensive inventory of dung-associated microbial taxa across all sampling months rather than investigating the dung samples separately.

To identify dung-associated taxa, we first extracted ASVs present in at least two of the 15 dung samples to build a reference database of dung-associated sequences. Gut-specific ASVs were then compared against this database using BLASTN. Gut ASVs exhibiting >99% sequence identity and >99% alignment coverage to dung reference sequences were classified as dung-associated and removed from fly microbiome datasets, generating a reduced “dung-filtered” dataset. All diversity analyses, ordinations, and indicator-species analyses were repeated using both the complete and dung-filtered datasets.

To quantify overlap between fly and dung microbiomes, we calculated for each fly sample (i) the frequency-weighted overlap (proportion of rarefied read counts attributable to dung-associated ASVs), and (ii) the absolute overlap (proportion of distinct ASVs shared with the dung-derived reference). Binomial mixed-effects models (GLMM with sample as random intercept), implemented via the *afex* (Singmann *et al*., 2025) and *lme4* (Bates *et al*., 2015) packages in R, were used to test whether Species and Season explained variation in the proportion of dung-associated reads across samples.

To assess the degree of taxon sharing among host species, we performed multi-species overlap analyses using the SuperExactTest framework (Wang *et al*., 2015) implemented in the *SuperExactTest* R package, separately for the complete dataset and for the dung-filtered dataset. Analyses were carried out at three taxonomic levels: individual ASVs, genus, and family. For each level, we first summed all read counts per taxon across all samples belonging to each host species, generating a species-level taxon list weighted by total read abundance. To control for differences in total read depth among species (arising from unequal sample sizes), we additionally performed a rarefied version of the analysis in which the species-level taxon list was randomly resampled (without replacement, seed = 20) to the size of the least-abundant species before testing. The background was defined as the total number of taxa observed across all samples at that taxonomic level (469 genera, or 10,388 / 10,334 ASVs for full and dung-filtered datasets, respectively). For each possible species combination (2^6^ - 1 intersections), the SuperExactTest framework calculated the observed overlap, expected overlap under a random hypergeometric model, fold-enrichment (FE = observed/expected), and an exact *P*-value.

## Results

### Seasonal Sampling

During the first year of the COVID-19 pandemic (2020), we repeatedly sampled a meadow at Ziegelhütte in Schwamendingen, Zurich, Switzerland, across 12 consecutive time points from April to Oktober. In total, we collected 4,685 male specimens representing 10 species of the dung fly genus *Sepsis*, all of which could be reliably identified to species level (Figure 1). The assemblage was strongly dominated by *Sepsis cynipsea*, which accounted for 80% of all collected individuals (3,751 specimens), while *S. punctum* was the rarest species represented by only eight individuals (0.1% of all specimens) throughout the entire sampled season. Consistent with a previous study (Rohner, Haenni, *et al*., 2019), species composition varied markedly over the course of the season. Whereas *S. neocynipsea* and *S. violacea* were most abundant during spring, *S. thoracica* and, to a lesser extent, *S. flavimana* increased in abundance toward the end of the sampling period.

**Figure 1.**
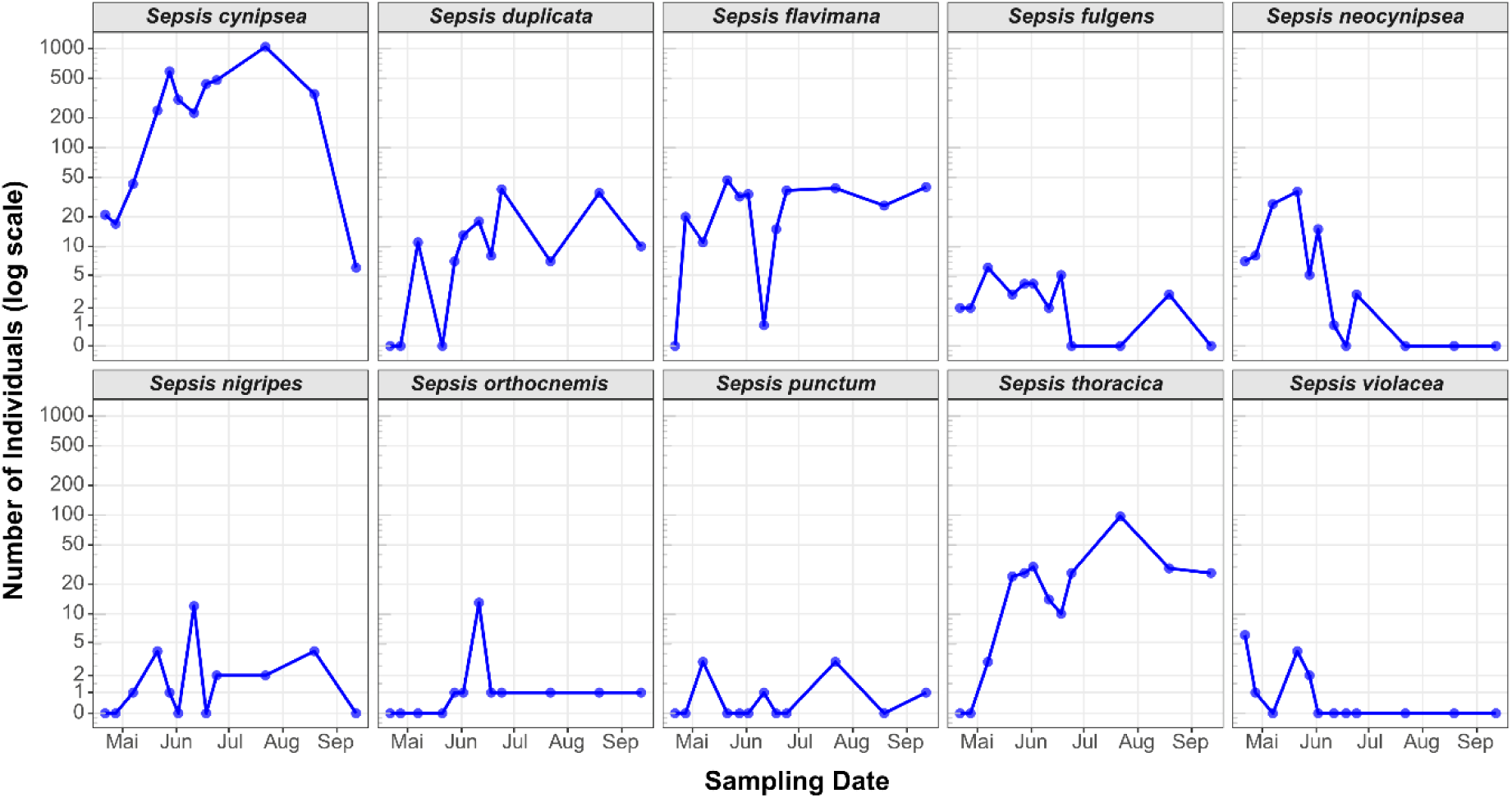
Seasonal dynamics of *Sepsis* species abundance at Ziegelhütte, Switzerland, during the 2020 sampling season. Abundances of ten *Sepsis* species collected from a meadow in Schwamendingen, Zurich, Switzerland, are shown across 12 consecutive sampling events between April and September 2020. Each panel represents a single species, with points connected by lines to illustrate temporal changes in abundance. The y-axis is displayed on a pseudo-logarithmic scale (including zeros) to facilitate comparison among species that differed markedly in abundance. *Sepsis cynipsea* was by far the most abundant species throughout the season, whereas several species, including *S. punctum*, *S. orthocnemis*, and *S. nigripes*, were encountered only sporadically in low numbers. Seasonal shifts in community composition were evident, with species such as *S. neocynipsea* and *S. violacea* occurring primarily during spring, while *S. thoracica* reached peak abundance later in the season.

To investigate species- and season-specific variation in the gut microbiome, we focused on six *Sepsis* species that were sufficiently abundant to permit repeated sampling across the active season. Individuals were collected at six time points between April and September 2020, providing temporal coverage of the major phases of seasonal community turnover (Rohner *et al*., 2015; Rohner, Haenni, *et al*., 2019). In total, 76 flies were selected for 16S rRNA amplicon sequencing, with three to four individuals per species and sampling date whenever possible (Table S2). The only exception was *S. duplicata*, which was recovered in sufficient numbers during only one of the six sampling events and is therefore represented by a single sampling date only. In addition, 15 dung samples collected across the season were sequenced to characterize the environmental microbial community and identify bacterial taxa likely originating from the dung substrate rather than the host gut microbiome.

### Sequencing Performance and Data Processing

PacBio sequencing generated a total of 3,099,186 circular consensus sequence (CCS) reads across 96 libraries. Following primer removal, quality filtering, denoising, and chimera detection, we retained 1,686,334 high-quality non-chimeric reads from 74 sepsid fly samples. Retention rates were generally high for fly samples, with a median retention of 74.7% relative to initial CCS reads. Two samples (Scyn_Apr_2 and Sfla_Apr_2) had to be excluded from subsequent analyses since they only yielded 1,051 and 393 reads, respectively, after rigorous filtering (see Table S3).

Rarefaction analyses indicated that most samples reached asymptotic diversity estimates at sequencing depths between 1,000 and 5,000 reads, suggesting that sequencing effort was sufficient to recover the majority of abundant bacterial taxa. Therefore, to ensure comparability among samples and reduce biases associated with unequal sequencing depth, we performed all subsequent analyses on datasets rarefied to 1,392 reads per sample. The complete sepsid dataset comprised 74 samples and 10,388 bacterial features (ASVs; Table S4).

In contrast, dung samples exhibited substantial read loss during sequence processing. Of the 297,358 CCS reads obtained from 15 dung samples, only 15,275 reads remained after denoising and chimera removal, corresponding to a median retention rate of 4.2%. Inspection of the retained reads revealed highly heterogeneous sequence compositions, suggesting an exceptionally diverse microbial community with many low-abundance sequence variants. Under these conditions, the DADA2 algorithm may have been unable to reliably reconstruct amplicon sequence variants (ASVs), as many true variants were represented by too few reads to be distinguished from sequencing errors. As a result, only 126 ASVs were detected across all dung samples (Table S5). We therefore refrained from conducting an in-depth analysis of the dung microbiome and instead retained only ASVs detected in at least two dung samples to distinguish environmental from host-associated microbes. Nevertheless, we caution that this dataset is likely biased toward the most abundant environmental taxa, potentially leading to an underestimation of the true contribution of environmental microbes to the sepsid gut microbiome.

### Overall Composition of the Sepsid Gut Microbiome

Across all host species and sampling periods, gut microbiomes were dominated by a relatively small number of bacterial genera despite the high total ASV richness.

The most abundant genera in the full dataset were *Commensalibacter* (23.92% of all reads), *Acinetobacter* (14.01%), *Romboutsia* (9.57%), *Pseudomonas* (6.56%), and *Enterococcus* (5.00%). Together, these genera accounted for more than half of all bacterial reads recovered from sepsid flies (Figure 2A).

**Figure 2.**
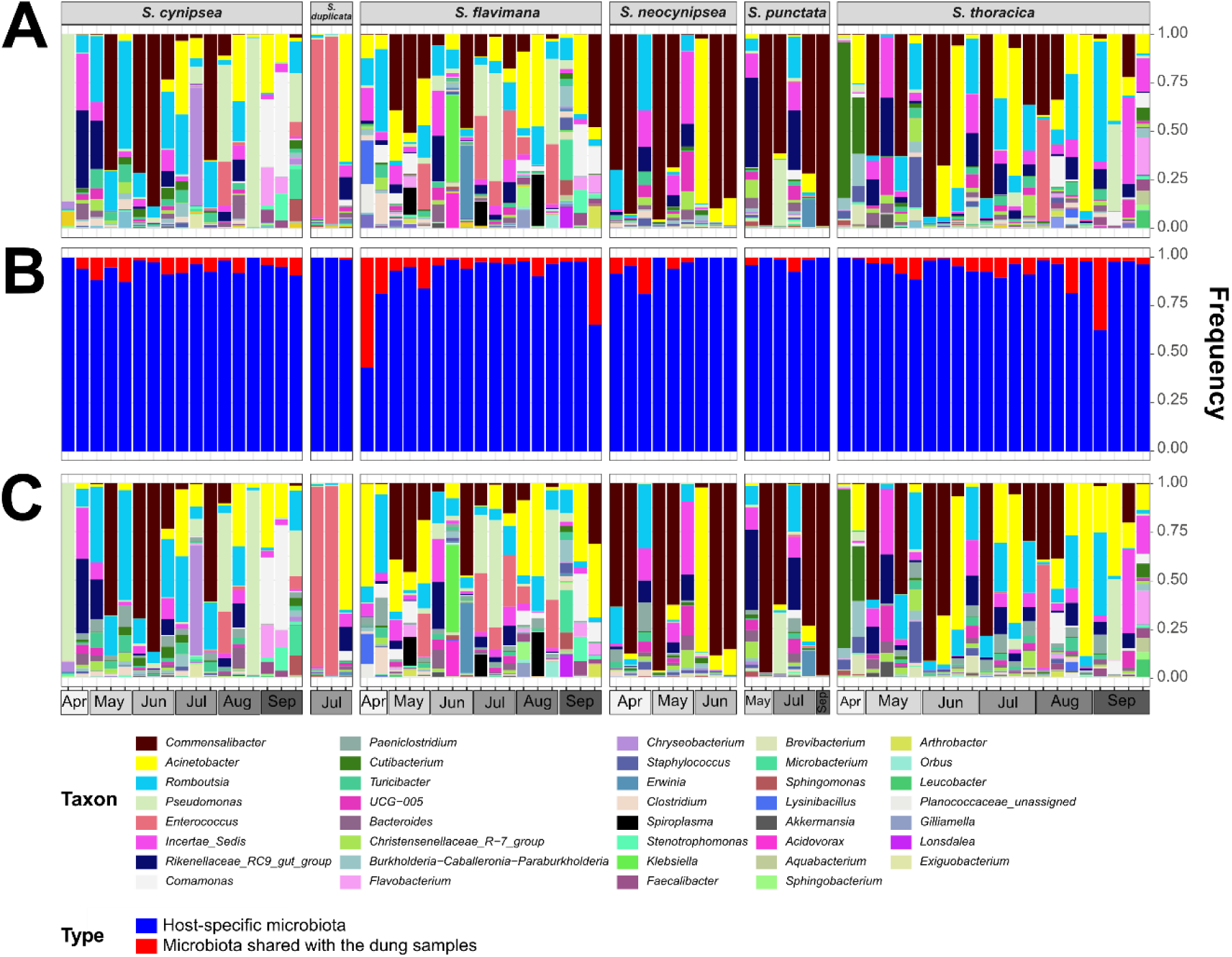
Seasonal variation in the gut microbiome composition of six sepsid fly species. Relative abundances of bacterial taxa are shown for individual samples of *Sepsis cynipsea*, *S. flavimana*, *S. neocynipsea*, *S. punctum*, and *S. thoracica* collected between April and September. Samples are grouped by host species and ordered chronologically by sampling month. In panel A, stacked bars depict the complete gut microbiome composition of each individual. The middle panel B shows the proportion of the microbiome classified as host-specific (blue) and the proportion shared with dung samples (red). Dung-associated taxa were defined as taxa occurring in at least two dung samples. In the lower panel C, these dung-associated taxa were removed, revealing the remaining host-specific microbiome composition. Colors denote bacterial taxa and are consistent between the upper and lower panels to facilitate comparison. Only taxa exceeding 5% relative abundance in at least one sample are displayed; lower-abundance taxa were omitted for clarity. Relative abundances are expressed as read count proportions of the total microbial community within each sample.

Substantial variation in taxonomic composition was observed among individual samples and across seasons. At the phylum level, communities were dominated by Proteobacteria and Firmicutes, while Bacteroidota and Actinobacteriota occurred at lower frequencies. Taxonomic profiles varied considerably through time, suggesting strong environmental influences on community assembly.

To determine the extent to which fly microbiomes reflected environmental microbial communities, we quantified the overlap between dung-associated and fly gut ASV communities. When dung-associated ASVs (those present in ≥2 dung samples) were used as a reference, an average of 6.3% of rarefied gut read counts per sample could be attributed to dung-associated taxa (frequency-weighted overlap; Figure 2B). The absolute overlap (proportion of distinct ASVs shared with dung) averaged 2.9% per sample. Overlap varied substantially among individual samples: some samples showed near-zero dung-associated reads, whereas others had dung-associated reads comprising a sizable minority of their microbiome (Figure 2B).

Binomial mixed-effects models revealed that host species, but not season, was a significant predictor of frequency-weighted dung overlap across all six species (χ^2^_5_ = 20.47, *P* = 0.001). In pairwise contrasts, *S. flavimana* showed significantly higher dung overlap than *S. duplicata* (log-odds ratio: −3.24, *z* = −3.27, *P* = 0.014), and *S. flavimana* also tended toward higher overlap than *S. punctum* (*P* = 0.012). When the analysis was restricted to the three species with the broadest seasonal coverage (*S. cynipsea*, *S. flavimana*, *S. thoracica*), a significant Species x Season interaction emerged (χ^2^_10_ = 29.72, *P* < 0.001), indicating that the seasonal pattern of environmental exposure differed among species.

Removal of dung-associated taxa altered the relative abundance of dominant taxa. Following removal of ASVs matching the pooled dung reference, *Acinetobacter* increased to 18.93% of total abundance and *Romboutsia* (15.15%), *Pseudomonas* (9.35%) as well as *Enterococcus* (7.0%) remained the dominant members of the residual community. These shifts indicate that environmentally acquired microbes and persistent host-associated microbes contribute variably to overall microbiome composition (Figure 2C).

In summary, these results suggest that gut microbiomes of sepsid flies contain a relatively small but species-dependent fraction of dung-associated microorganisms, and that the degree of environmental contamination differs among host species rather than uniformly with season.

### The Core Microbiome

Removal of dung-associated taxa revealed a smaller but highly structured host-associated microbiome. Redundancy analyses of the dung-filtered dataset continued to identify significant effects of both host species and season at the genus level, indicating that ecological structure persisted even after environmental taxa were removed. Core microbiome indicator analyses identified the same set of host-species and seasonal indicators as in the full dataset, including *Commensalibacter* as a persistent host-associated taxon and *Acinetobacter* and *Comamonas* as seasonal associates.

Multi-species taxon-sharing analyses revealed extreme and highly significant enrichment of shared taxa across all six fly species at every taxonomic level examined (Table S6). In the complete dataset, read-depth-equalised (i.e. rarified) genus-level analyses identified 36 genera shared by all six *Sepsis* species (background: 469 genera; FE = 119.3; *P* = 2.32 x 10^-71^). Without rarefaction, 52 genera were shared (FE = 20.7; *P* = 1.83 x 10^-62^). At the family level, 49 families were shared by all six species (background: 195 families; FE = 8.8; *P* = 5.70 x 10^-48^). At the ASV level, 19 sequence variants (rarified) or 25 (non-rarified) were shared across all six species (rarified: FE = 6,711; *P* = 9.39 x 10^-67^). Five-species subsets consistently showed 33- to 46-fold enrichment above random expectation (all *P* < 10^-5^).

In the dung-filtered dataset, results were comparable or even stronger. Rarified genus-level analyses identified 38 genera (FE = 120.0; *P* = 6.79 x 10^-76^) and non-rarified analyses 47 genera (FE = 19.9; *P* = 1.13 x 10^-54^) shared by all six species. At the family level, 49 families were shared (FE = 9.6; *P* = 2.34 x 10^-50^). At the ASV level, 16 (rarified) or 22 (non-rarified) ASVs were shared (rarified: FE = 6,680; *P* = 2.31 x 10^-56^).

Comparing the full and rarified dung-filtered genus-level cores, 31 genera were present in both core sets, constituting a robust core that persists regardless of environmental contamination. These 31 genera include *Acinetobacter*, *Romboutsia*, *Pseudomonas*, *Alistipes, Bacteroides, Christensenellaceae R-7 group, Cutibacterium, Enhydrobacter, Herbaspirillum, Methylobacterium, Paracoccus, Phascolarctobacterium, Sphingomonas, Stenotrophomonas, Staphylococcus, Turicibacter*, and Rikenellaceae RC9 gut group, among others. Five genera present only in the full core (*Brevundimonas*, *Candidatus Saccharimonas, Paeniclostridium*, Prevotellaceae UCG-001, Prevotellaceae UCG-003) may overlap with dung communities and thus dropped out after dung filtering. Seven genera newly appearing in the dung-filtered core (*Acetitomaculum*, *Aquabacterium, Comamonas, Parasutterella, Erwinia,* Erysipelotrichaceae UCG-009, Candidatus Soleaferrea) are likely genuine host-associated taxa that were partially masked in the full dataset by the presence of dung-associated reads. These results demonstrate that sepsid flies contain a large and highly non-random host-associated core microbiome community that persists despite substantial seasonal turnover and continual exposure to environmental microbes, even after statistical removal of dung-associated taxa.

### Alpha Diversity

Alpha diversity analyses revealed contrasting patterns depending on the diversity metric used, both in the dung-filtered (presented here and in Table S7) and the full dataset (see Table S7).

Bacterial Shannon diversity did not differ significantly among sepsid host species, sampling months, or their interaction (Species: *F*_5,49_ = 1.4, *P* = 0.239; Season: *F*_5,49_ = 1.1, *P* = 0.376). Because the Shannon index integrates both richness and evenness, this result indicates that overall bacterial diversity remained broadly similar across species and seasons. Similarly, analyses of Chao1 richness (Species: *F*_5,49_ = 1.45, *P* = 0.22; Season: *F*_5,49_ = 0.36, *P* = 0.872) and observed-feature richness (Species: *F*_5,49_ = 0.72, *P* = 0.613; Season: *F*_5,49_ = 0.67, *P* = 0.645) revealed no significant differences, confirming that microbiome taxon richness did not vary systematically among host fly species or seasons.

By contrast, Simpson bacterial diversity differed significantly among host fly Species (*F*_5,49_ = 4.36, *P* = 0.002), whereas Season or the Species-by-Season interaction did not. Pielou’s evenness (J’) also showed a significant Species effect (*F*_5,49_ = 2.51, *P* = 0.042), indicating that species differences are driven by community evenness rather than richness.

Analysis of the full (i.e. unfiltered) dataset yielded quantitatively similar results, confirming no seasonal or species-specific patterns of alpha diversity on the transient microbiome. Taken together, alpha-diversity analyses indicate that host species harbor comparable numbers of bacterial taxa but differ somewhat in the relative abundance distributions of those taxa, with some fly species harboring more even and others more uneven microbial communities.

### Beta Diversity

Principal coordinate analyses based on Bray-Curtis dissimilarity and weighted UniFrac distance revealed substantial variation among samples with partial clustering by season, both in the dung-filtered (presented here and in Table S8) and the full dataset (see Table S8; Figure 3).

**Figure 3.**
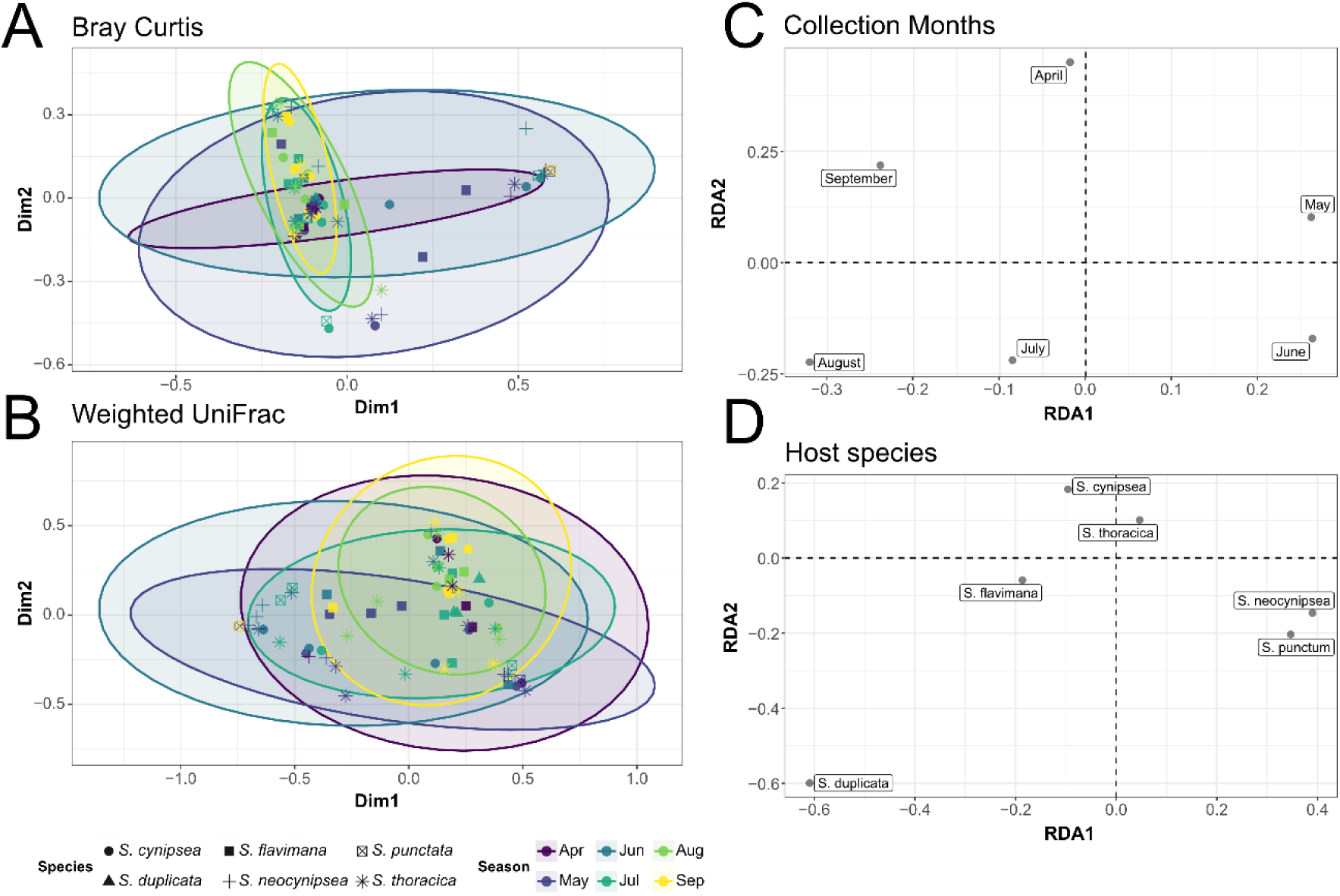
Seasonal and host-species variation in gut microbiome composition. (A,B) Principal coordinate analyses (PCoA) based on beta-diversity distances among individual gut microbiome samples. Points represent individual flies, with colors indicating sampling month and symbols denoting host species. Ellipses encompass seasonal variation in ordination space. (C) Redundancy analysis (RDA) showing the relationship between gut microbiome composition and sampling month. Sampling months are displayed as centroids in ordination space illustrating seasonal shifts in microbiome structure. (D) Redundancy analysis (RDA) showing differences in gut microbiome composition among host species. Species centroids indicate taxon-specific microbiome signatures after accounting for variation in community composition. Together, the ordinations reveal both seasonal and host-species effects on gut microbiome composition, with *S. duplicata* showing the strongest separation from the remaining species, and spring samples occupying a distinct region of ordination space relative to summer and autumn samples.

Bray-Curtis PERMANOVA analyses further detected significant pairwise differences among several seasonal comparisons (Figure 3A). After Benjamini-Hochberg FDR correction, significantly distinct communities were detected between April and August (*q* = 0.045) as well as between May and September (*q* = 0.045). Additional comparisons - including May vs.

August (*q* = 0.07), June vs. July (*q* = 0.090), May vs. July (*q* = 0.090), and June vs. September (*q* = 0.090) - differed only marginally. These results indicate progressive temporal turnover in bacterial microbiome community composition over the growing season, particularly contrasting spring and summer assemblages. Curiously, we did not find significant differences between the earliest sampling time point in April and the latest in September (*q* = 0.093). In contrast to season, our Bray-Curtis analyses only detected marginal species-level differences, with *S. flavimana* differing from both *S. neocynipsea* (*q* = 0.055) and *S. punctum* (*q* = 0.055), and *S. punctum* differing from *S. thoracica* (*q* = 0.055).

Weighted UniFrac analyses failed to detect significant pairwise differences among seasonal samples or host species after FDR correction, with all *q*-values substantially exceeding 0.05 (Figure 3B).

For the full dataset, we found overall lower *q*-values, suggesting that subsetting and removal of microbes lowered our statistical power. The discrepancy between Bray-Curtis and weighted UniFrac results may indicate that seasonal and species-level turnover primarily involved replacement among closely related bacterial taxa rather than turnover among phylogenetically distinct lineages. Thus, community composition changed substantially over the season while broader phylogenetic structure remained comparatively stable.

### Seasonal and Host Effects on Microbiome Community Composition

Redundancy analyses (RDA) identified both host species and season as significant determinants of microbiome composition across all taxonomic levels examined both for the dung-filtered (shown here) and the full microbial dataset (see Table S9). In the complete dataset at the genus level, both Season (*F*_5,63_ = 1.85, *P* = 0.003) and Species (*F*_5,63_ = 1.75, *P* = 0.007) explained significant variation (Type III permutation tests).

Forward model selection entered Season first (adjusted *R*^2^ = 0.064), followed by Species (full model adjusted *R*^2^ = 0.113), indicating that season consistently explained a larger proportion of variance than host species. Results were qualitatively similar at the species level (Season: *F*_5,63_ = 1.96, *P* = 0.001; Species: *F*_5,63_ = 1.72, *P* = 0.005) and other taxonomic levels. Corresponding ordination plots revealed clear separation of communities along seasonal gradients, particularly contrasting spring and late-summer samples. Host species also exhibited significant clustering, although species-specific patterns were generally weaker than seasonal effects (Figure 3C and D).

Analyses of the dung-filtered dataset retained significant effects of both Season (*F*_5,63_ = 1.80, *P* = 0.004) and Species (*F*_5,63_ = 1.66, *P* = 0.006) at the genus level (full model adjusted *R*^2^ = 0.105), demonstrating that host-species effects on microbiome composition are not attributable solely to differences in dung-microbe exposure. Collectively, these findings indicate that seasonal environmental variation is the dominant driver of microbiome composition, while host species exert a secondary but persistent filtering effect.

### Bacterial Indicator Taxa Associated with Host Species and Season

Indicator-species analyses identified bacterial taxa associated with particular host species and sampling periods. At the genus level, host-associated indicator taxa included *Pantoea* for *S. duplicata* (stat = 0.684, *P* = 0.021) and *Aquabacterium* for *S. thoracica* (stat = 0.785, *P* = 0.007). *Enterococcus* was significantly associated with a grouping comprising *S. cynipsea*, *S. duplicata*, and *S. flavimana* (stat = 0.773, *P* = 0.026); *Brevibacterium* with a grouping of *S. flavimana*, *S. neocynipsea*, *S. punctum*, and *S. thoracica* (stat = 0.804, *P* = 0.009); and *Commensalibacter* with a broad grouping comprising all species except *S. duplicata* (stat = 0.880, *P* = 0.029). At the species level, *S. duplicata* additionally showed significant association with *Enterococcus wangshanyuanii* (stat = 0.777, *P* = 0.002), *Acinetobacter piscicola* (stat = 0.728, *P* = 0.012), and *Pantoea agglomerans* (stat = 0.630, *P* = 0.032).

Seasonal associations were considerably more common, with 15 significant indicator taxa identified at the genus level. *Bacillus* was strongly associated with May samples (stat = 0.791, *P* < 0.001). Communities spanning July and August showed elevated *Enterococcus* (stat = 0.823, *P* < 0.001). Late-season communities (August-September) were characterized by *Comamonas* (stat = 0.783, *P* < 0.001) and *Arthrobacter* (stat = 0.678, P = 0.003).

September samples showed elevated *Flavobacterium* (stat = 0.785, *P* < 0.001), *Faecalibacter* (stat = 0.781, *P* < 0.001), and *Leucobacter* (stat = 0.688, *P* = 0.002). September communities were specifically characterized by *Stenotrophomonas ginsengisoli* and *Acinetobacter radioresistens*. Early-season communities (April-May) were associated with *Mogibacterium* and the Family XIII AD3011 group. Communities spanning mid-to-late season (July through September) were associated with *Microbacterium*, while Christensenellaceae R-7 group and *Olsenella* were associated with early-to-mid season (April through July) samples.

The greater number of seasonal than host-associated indicator taxa further support the conclusion that temporal environmental variation represents the primary source of microbiome turnover. Results from the dung-filtered dataset produced a qualitatively similar indicator set, confirming that these associations are not simply a consequence of differential dung-microbe exposure.

### No evidence for *S. thoracica* male morph-specific microbiome differences

The two distinct male morphotypes of *S. thoracica* that substantially differ in size and coloration (Busso *et al*., 2017; Busso and Blanckenhorn, 2018b, 2018a; Blanckenhorn *et al*., 2020; Gourgoulianni *et al*., 2023) do not systematically differ in their gut microbiome composition. Redundancy analyses of their microbiome across species-, genus-, family-, and order-level consistently showed that body size explained only a small proportion of community variation and was never significantly associated with overall microbiome structure, whereas seasonal effects explained a larger fraction of the variation and repeatedly showed weak trends towards significance (Table S10). Indicator species analyses identified several taxa associated with seasonal groups, but only one single bacterial lineage within the Staphylococcales that was consistently associated with morph size across taxonomic levels. These results indicate that gut microbial communities are largely conserved in both male morphs, with any size-related differences being limited to a very small subset of microbial taxa rather than reflecting broad changes in community composition. These findings were consistent for the full and the dung-filtered dataset.

## Discussion

Our study provides the first comprehensive characterizations of gut microbiome variation in sepsid dung flies and demonstrates that microbiome assembly in these insects is jointly shaped by seasonal environmental variation and host-associated filtering. By combining environmental dung samples with seasonal sampling and multiple ecological partitions of the data, we were able to separate environmentally inherited microbial variation from host-associated community structure. Across all analyses, seasonal variation emerged as the strongest and most consistent determinant of microbiome composition, whereas host species exerted weaker but persistent effects. At the same time, a significant core microbiome remained detectable after removal of dung-associated taxa, indicating that sepsid gut communities are not simply transient reflections of environmental microbial exposure.

### Seasonal Environmental Filtering Dominates Dung Fly Microbiome Assembly

The most consistent pattern across all analyses was the strong effect of season on dung fly microbiome composition. Seasonal effects were detected in redundancy analyses across nearly all taxonomic levels and remained significant after removal of environmentally associated taxa. Moreover, indicator-species analyses identified substantially more season-associated than host-associated taxa, suggesting that temporal environmental variation is the dominant source of microbiome turnover.

This result is not surprising given the ecology of dung habitats (Floate, 2023). Fresh cow dung represents a highly dynamic microbial environment that undergoes rapid successional changes driven by temperature, moisture, oxygen availability, nutrient depletion, and microbial interactions (Holter, 2016). Because sepsid flies feed, mate, and oviposit directly on dung, they are continuously exposed to these changing microbial communities. Consequently, temporal turnover in environmental microbial communities is expected to generate corresponding changes in the microorganisms available for gut colonization.

The importance of environmental acquisition of microbes observed here is consistent with growing evidence that many insect microbiomes are assembled primarily through repeated environmental recruitment rather than through stable vertical transmission (Hammer *et al*., 2019). In contrast to obligate symbioses such as those found in aphids, carpenter ants, or tsetse flies, microbiomes of many holometabolous insects appear highly dynamic and strongly influenced by ecological context (Engel and Moran, 2013; Douglas, 2015). Our results demonstrate that sepsid dung flies belong largely to this environmentally acquired end of the insect microbiome spectrum.

The contrast between Bray-Curtis and weighted UniFrac analyses provides further insight into this context. Significant differences among early and late-season microbial communities were detected using Bray-Curtis dissimilarity but not by weighted UniFrac. As Bray-Curtis considers taxon abundances and UniFrac additionally incorporates phylogenetic relationships, this discrepancy suggests that seasonal turnover primarily involves replacement of closely related bacterial taxa rather than replacement of major phylogenetic lineages. Thus, community composition changes substantially over time while broader phylogenetic structure remains comparatively stable. Curiously, the earliest (April) and latest (September) sampling months did not differ significantly in gut microbiome composition, whereas other spring–autumn comparisons, such as April-August and May-September, were significantly different. This pattern suggests that seasonal changes in the microbiome are not strictly directional throughout the year. Instead, the similarity between spring and autumn communities may indicate that microbiome dynamics follow a cyclical seasonal trajectory, with communities at the beginning and end of the active season converging on a similar compositional state. This hypothesis is further supported by the circular distribution of sampling months along the first two RDA axes (Figure 3C), which suggests a recurring annual cycle in community composition rather than a linear temporal progression.

Several non-exclusive mechanisms could explain this pattern. First, components of the microbiome may persist through adult fly overwintering (see Zeender *et al*., 2019) and influence the composition of spring communities. Consistent with this idea, Ferguson et al. (2018) showed that gut microbial communities of the spring field cricket *Gryllus veletis* undergo pronounced changes during overwintering that are closely associated with shifts in cold tolerance and immune function. Second, flies may reacquire microbes from environmental reservoirs which themselves persist across seasons. Because many insect-associated bacteria are environmentally acquired, microbial communities present in dung, soil, or other overwintering substrates may act as source pools from which flies are recolonized each spring. Given that dung pats are ephemeral by nature and disintegrate completely with time and over winter, this mechanism seems less plausible however. Third, seasonal environmental conditions may repeatedly favor similar microbial assemblages at the beginning and end of the active season, as, for example, found in mouse populations (Marsh *et al*., 2022). Under this scenario, comparable temperature regimes, resource availability, or host physiological states in spring and autumn could generate similar microbiome compositions despite the temporal separation between sampling periods.

Seasonal restructuring of insect-associated microbiomes has been documented in several other species. In honey bees (*Apis mellifera*), gut microbiome composition differs markedly between winter and summer workers, with winter bees harboring distinct bacterial communities and altered bacterial loads than summer nurses and foragers (Kešnerová *et al*., 2020; Castelli *et al*., 2022). Similarly, overwintering-associated changes in gut microbiome composition have been reported in the spring field cricket *G. veletis*, where microbiome shifts occur concurrently with physiological adaptations to winter conditions (Ferguson *et al*., 2018). Together, these studies suggest that seasonal transitions can strongly influence insect microbiomes and provide a plausible framework for interpreting the unexpected similarity between spring and autumn microbiomes observed in *Sepsis*.

However, distinguishing between microbiome persistence through overwintering, recolonization from environmental reservoirs, and cyclical responses to seasonal environmental conditions will require targeted sampling across multiple years and overwintering stages (cf. Zeender *et al*., 2019).

### Host Species Impose Secondary Filtering on Environmental Communities

Despite the dominant role of season demonstrated here, host species additionally define microbiome composition. Importantly, species differences persisted after removal of dung- associated microbial taxa and remained detectable across multiple analytical approaches. These findings indicate that microbiome composition cannot be explained solely by environmental exposure.

Host-associated filtering could arise through several mechanisms. Species may differ in gut morphology, digestive physiology, immune responses, developmental trajectories, or behaviour, all of which can influence microbial persistence within the gut. Even subtle differences in feeding behaviour or microhabitat use may alter exposure to environmental microbial communities. Because several *Sepsis* species coexist within the same dung pats while exhibiting differences in phenology and life-history traits (Rohner, Roy, *et al*., 2019; Blanckenhorn *et al*., 2020, 2021), species-specific filtering is biologically plausible.

Interestingly, alpha-diversity analyses indicated that species differed in community evenness but not richness. Shannon diversity did not vary significantly among species, whereas Simpson diversity did. This pattern suggests that host species do not differ substantially in the total number of microbial taxa they harbor. Instead, species-specific effects appear to arise through differential enrichment of particular taxa. Such changes in relative abundances of microbiomes are consistent with host filtering mechanisms that influence microbial persistence rather than colonization itself.

Although host-species effects were weaker than seasonal effects, they remained remarkably robust across multiple sensitivity analyses. This persistence supports the view that host-associated processes contribute meaningfully to microbiome assembly even in insects that experience continuous exposure to environmentally derived microorganisms, as is the case in dung flies and beetles (Skidmore, 1991; Floate, 2011, 2023; Hanski and Cambefort, 2014).

### A Large, Persistent and Genuinely Host-Associated Core Microbiome

A particularly striking finding of this study is the extreme degree of non-random genus sharing among all six sympatric *Sepsis* species. In read-depth-equalised analyses, 36 genera were shared by all six species in the full dataset, representing ∼119-fold enrichment above random expectation, far exceeding what would be expected from stochastic co-occurrence in a shared habitat. Crucially, this core was not merely a reflection of a shared environmental source pool: after removal of dung-associated taxa, 38 genera were shared at comparable enrichment, and 31 genera were present in both the full and dung-filtered cores. These results strongly imply that the core microbiome represents biologically meaningful, host-associated bacterial communities rather than transiently ingested environmental organisms. There likely is a strong phylogenetic signal, as European *Sepsis* flies are closely related (Ang *et al*., 2013; Zhao *et al*., 2013).

The existence of such a large and highly enriched core is notable because it emerged despite extensive environmental exposure and strong seasonal turnover. If gut communities merely reflected random subsets of dung microorganisms, removal of dung-associated taxa should have largely eliminated detectable sharing structure. Instead, after dung filtering 38 genera remained shared at 120-fold enrichment, similar to the full dataset results. These findings support a two-layer model of microbiome assembly: a transient environmentally acquired community is superimposed on a more stable, host-associated core that persists across species, seasons, and environmental conditions.

Five genera present only in the full core (*Paeniclostridium, Prevotellaceae UCG-001/003, Brevundimonas, Candidatus, Saccharimonas*) were not detected in the dung-filtered core, suggesting they stem from dung communities and represent conditionally shared environmental taxa. Conversely, seven genera appeared in the dung-filtered core but not the full core (including *Comamonas*, *Aquabacterium*, *Parasutterella*, *Acetitomaculum*), which may represent genuine host-associated taxa whose signal was partially obscured by co-occurring dung-associated reads in the full dataset. The 31 genera consistently shared in both analyses - including *Acinetobacter*, *Romboutsia*, *Pseudomonas*, *Alistipes*, *Bacteroides*, Christensenellaceae R-7 group, *Cutibacterium*, *Herbaspirillum*, *Sphingomonas*, *Staphylococcus, Stenotrophomonas,* and *Turicibacter* - represent the most robust candidates for genuinely host-associated bacterial genera of sepsid flies.

Similar patterns have been reported in drosophilids, where a small but recurrent set of bacterial taxa persists across hosts and environments despite substantial temporal turnover (Wong *et al*., 2011, 2015; Pais *et al*., 2018; Wang *et al*., 2020). Our results suggest that a comparable process operates in sepsid flies, but with a notably larger core: 36-38 shared genera and 16-19 shared ASVs represent substantial host-associated diversity. The enrichment factors (∼120-fold at genus level; ∼6,700-fold at ASV level) are exceptionally high when compared to other multi-species insect microbiome studies, suggesting that *Sepsis* species maintain particularly consistent gut microbial associations despite their generalist coprophagous lifestyle.

### Ecological Significance of *Commensalibacter*

Among all microbial taxa recovered in this study, *Commensalibacter* is particularly noteworthy. This genus represented the most abundant bacterial lineage in the full dataset and remained highly abundant after removal of dung-associated taxa. Furthermore, *Commensalibacter* repeatedly emerged as an indicator taxon within host-associated community analyses.

Members of the genus *Commensalibacter* belong to the family Acetobacteraceae and are common associates of drosophilid flies and other insects (Botero *et al*., 2023). Experimental work in *Drosophila* has demonstrated that Acetobacteraceae can influence host nutrition, growth, and gut homeostasis (Pais *et al*., 2018) and contributes to immune regulation in honey bees (Siozios *et al*., 2019). Several species are capable of persistent gut colonization and may exhibit greater host association than many environmentally acquired taxa (Botero *et al*., 2023).

The repeated occurrence of *Commensalibacter* across wild populations of *Drosophila* (see, for example, Kapun *et al*., 2020) and multiple *Sepsis* species suggests that some bacterial associations may be conserved across higher Diptera. Although the functional significance of this genus in sepsids remains unknown, its prevalence and persistence make it a promising target for future experimental work. In particular, studies manipulating *Commensalibacter* abundance could help determine whether this taxon contributes to nutrition, digestion, pathogen resistance, or any other significant aspect of host biology.

### Gut-Dung Overlap is Low and Species-Dependent

Comparisons between dung and gut microbiomes revealed that merely a small fraction of gut read abundance (on average 6.3% per sample by frequency weighting) could be attributed to ASVs also detected in the pooled dung community. This low overlap indicates that the great majority of gut microbiome sequences represent taxa not detectable in the broader dung environment, consistent with substantial host-associated filtering or colonisation from sources other than the immediate dung environment.

Importantly, the proportion of dung-associated reads in the gut differed significantly among host species, with *S. flavimana* showing considerably higher environmental contamination than *S. duplicata*. Season was not a significant predictor of overlap in the full dataset, though a significant Species x Season interaction was detected when analyses were restricted to the most common three fly species, suggesting that the relationship between environmental exposure and gut colonisation is species-specific and context-dependent (Wang *et al*., 2020). This contrasts with a scenario in which gut microbiomes simply represent a passive reflection of the surrounding environmental microbial community. Instead, our results indicate that host-associated processes contribute to shaping gut microbial communities and may differ among host species.

However, these conclusions should be interpreted with caution. Our attempt to distinguish host-associated from environmentally derived microbes relied on identifying bacterial taxa present in dung samples. Following DADA2 processing and filtering, only a relatively small number of dung-associated ASVs remained available for downstream analyses. As a consequence, our ability to identify and remove environmentally acquired bacteria was limited and likely biased towards the most abundant and consistently detected taxa. Many rare taxa, as well as bacteria that were present only during specific sampling periods, may not have been detected in the dung dataset and therefore could not be classified as environmentally derived. Furthermore, because dung samples were pooled across sampling periods, we were unable to account for temporal variation in the environmental microbiome in this specific analysis. Consequently, some bacteria classified as host-associated may in fact represent environmentally acquired taxa that escaped detection in the dung samples.

Given these limitations, we refrain from overinterpreting patterns of host-specific microbiome assembly and instead view our results as preliminary evidence that host species identity contributes to shaping gut microbial communities beyond simple environmental exposure. Substantially deeper sequencing of the putatively hyper-diverse dung samples will be needed to get robust estimates of the microbiome diversity in dung samples.

### Implications for Ecological Coexistence and Niche Partitioning

Although the present study was not designed to test competition directly, the observed microbiome differences among sympatric species raise intriguing questions regarding ecological coexistence. Multiple *Sepsis* species coexist within the same habitat despite apparently similar life histories and larval diets, and such coexistence typically requires some degree of ecological differentiation (Rohner *et al*., 2015; Laux *et al*., 2019; Rohner, Haenni, *et al*., 2019; Zeender *et al*., 2019; Blanckenhorn *et al*., 2020, 2021).

One possibility is that microbiome differences reflect subtle differences in resource use. Species may exploit different stages of dung decomposition, occupy different microhabitats within dung pats, or preferentially consume distinct microbial resources. Because microorganisms constitute a major nutritional component of dung-based food webs, differences in microbiome composition could potentially reflect or contribute to niche differentiation (Shukla *et al*., 2016; Ebert *et al*., 2021).

Our results do not allow direct tests of these hypotheses. Nevertheless, the persistence of species-specific microbial signatures after removal of obvious environmental (i.e. dung) taxa is consistent with the possibility that host species interact differently with microbial resources. Future studies combining microbiome analyses with experimental manipulations of diet, resource availability, and microbial communities will be required to determine whether and how microbiome variation contributes directly to ecological coexistence.

### Limitations and Future Directions

Beyond the limitations associated with our dung samples discussed above, particularly the limited quantitative data that prevented a comprehensive characterization of the dung microbiome, several additional factors should be considered when interpreting our findings. First, our study was conducted at a single location during a single growing season. Although repeated temporal sampling provides strong evidence for seasonal effects, additional years and geographic locations will be necessary to evaluate the generality of observed patterns. Second, our analyses focused exclusively on adult males. Because microbiomes can vary across developmental stages and the sexes, future work should include larvae, pupae, and females. Third, full-length 16S sequencing provides high taxonomic resolution but does not directly measure microbial function. Functional predictions based on PICRUSt2 and trait databases yielded no strong metabolic signal, likely reflecting limited representation of dung-associated bacteria in existing reference databases. Metagenomic and metatranscriptomic approaches will therefore be necessary to determine whether taxonomic differences correspond to functional differences of microbial communities.

Importantly, experimental manipulations remain essential for establishing causal relationships between microbiomes and host ecology. Controlled inoculation experiments, microbiome transplants, or gnotobiotic rearing approaches could determine whether particular bacterial taxa influence host fitness, resource utilization, or competitive interactions.

## Conclusions

Our results demonstrate that microbiome assembly in sepsid dung flies is governed by the interaction of seasonal environmental filtering and host-associated selection. Seasonal turnover in environmental microbial communities provides the primary source of variation, while host species selectively retain a subset of taxa that generate persistent microbiome structure. Despite continual exposure to highly diverse dung-associated microorganisms, a significant core microbiome thus persists across species and seasons. These findings support a hierarchical model of microbiome assembly and suggest that host-microbe interactions may represent an underappreciated component of ecological differentiation in dung fly communities. More broadly, our study highlights the importance of integrating environmental and host-associated perspectives when investigating microbiome assembly in natural populations.

## Acknowledgments

We thank Fabian Staubach for his support and advice regarding statistical analyses, and Marcel Suleiman for his guidance and assistance with DNA extraction and amplicon generation. We also thank the Functional Genomics Center Zürich for sequencing services and the Department of Evolutionary Biology and Environmental Studies at the university Zurich for funding the PacBio sequencing. We acknowledge the use of GitHub Copilot (Claude Sonnet 4.6) for assistance with code generation, code auditing, and documentation, and ChatGPT (OpenAI) for assistance in improving the clarity and readability of the manuscript text.

## Data Availability

Raw PacBio sequencing reads are deposited at the Short Read Archive (SRA) under accession number https://www.ncbi.nlm.nih.gov/bioproject/PRJNA1481660. The complete data analysis pipeline, including all custom scripts, is available on GitHub at https://github.com/capoony/SepsidMicroBiome.

## Authors contribution

MK: Conceptualization, Methodology, Software, Formal analysis, Data Curation, Writing - Original Draft, Visualization, Project administration, Funding acquisition, JR: Conceptualization, Validation, Investigation, Resources, Data Curation, Writing - Review & Editing, WB: Conceptualization, Validation, Resources, Writing - Review & Editing

## Ethics Statement

All sampling procedures and data analyses were carried out in accordance with established ethical guidelines and standards of good scientific practice.

